# Experience-dependent sharp-wave ripple deficits in an Alzheimer’s disease mouse model

**DOI:** 10.1101/2025.06.09.658771

**Authors:** Paulina Schnur, Nicole Byron, Symeon Gerasimou, Alba Pascual Clavel, Valentin Rohner, Tommaso Patriarchi, Shuzo Sakata

## Abstract

Amyloid pathology is a hallmark of Alzheimer’s disease (AD). Hippocampal sharp-wave ripples (SWRs) play a role in memory consolidation and are impaired in various AD mouse models. However, it remains unclear how experience affects SWRs and how extrinsic signals contribute to SWR generation in AD. Here, by combining behavioral, *in vivo* electrophysiological and fiber photometry approaches, we show that an experience-dependent increase in hippocampal SWRs is disrupted in male and female 5xFAD mice. In wild-type mice, SWRs during non-rapid eye movement sleep (NREMS) increased after exploring a novel environment and the SWR rate gradually decreased with NREMS episodes and multiple behavioral sessions. However, 5xFAD mice did not show such experience-dependent SWR rate changes. A similar deficit was observed after a novel object recognition test. On the other hand, sleep spindles were intact in 5xFAD mice under all conditions. Since deficits in basal forebrain cholinergic neurons have been implicated in 5xFAD mice and SWRs are regulated by hippocampal cholinergic tone, we examined if hippocampal cholinergic signals could explain experience-dependent SWR deficits. Using fiber photometry and expressing a genetically encoded acetylcholine (ACh) sensor in the hippocampus, we found that ACh dynamics in the hippocampus were intact in 5xFAD mice across sleep-wake cycles, including NREMS, while we also found a negative correlation of infraslow cortical signal power dynamics with hippocampal ACh signals during NREMS regardless of genotypes. These results suggest that experience-dependent SWR deficits stem from non-cholinergic pathological changes.

## Introduction

Alzheimer’s disease (AD), the leading cause of dementia, is characterized by pathological hallmarks of amyloid plaques and tauopathy (Hardy and Higgins, 1992; Karran et al., 2011; Long and Holtzman, 2019; Scheltens et al., 2021). Immunotherapies targeting amyloid-β oligomers have shown promising results in humans (Sims et al., 2023; van Dyck et al., 2023). However, given the limited efficacy of these emerging treatments, diversifying treatment options and identifying various biomarkers are urgently needed.

Electrophysiological biomarkers of AD have long been recognized (Jeong, 2004; Nakamura et al., 2018; Ranasinghe et al., 2020; Byron et al., 2021; Meghdadi et al., 2021; Schoonhoven et al., 2022; Kudo et al., 2024). A slowing down of neural oscillation frequencies is a consistent finding in both AD and mild cognitive impairment (Jeong, 2004). These abnormalities have been targeted for non-invasive and invasive neuromodulation (Lozano and Lipsman, 2013; Chang et al., 2018; Luo et al., 2021; Chan et al., 2022). Among the various oscillations, hippocampal sharp-wave ripples (SWRs) have been less explored in AD.

A SWR comprises a sharp wave and ripples. A sharp wave is generated by recurrent excitation in the hippocampal CA3, resulting in large transmembrane currents in the apical dendrites of CA1 pyramidal cells via Shaffer collaterals and high-frequency synchronous firing of CA1 neurons as ripples (Sullivan et al., 2011). SWRs are conserved across mammalian species and typically occur during quiet wakefulness and non-rapid eye movement sleep (NREMS) (Buzsaki, 2015; Liu et al., 2022).

Since SWRs are accompanied by the reactivation of neural ensembles, they have been implicated in systems consolidation (Wilson and McNaughton, 1994; Diba and Buzsaki, 2007; Foster, 2017; Joo and Frank, 2018). Indeed, SWRs can causally influence memory (Girardeau et al., 2009; Jadhav et al., 2012; Fernandez-Ruiz et al., 2019; Chang et al., 2025). Abnormalities in SWRs have been reported in various AD mouse models (Ciupek et al., 2015; Gillespie et al., 2016; Nicole et al., 2016; Witton et al., 2016; Hollnagel et al., 2019; Jones et al., 2019; Jura et al., 2019; Benthem et al., 2020; Caccavano et al., 2020; Prince et al., 2021; Funane et al., 2022). Pathological SWRs in AD are, however, not fully understood.

Among various AD mouse models, 5xFAD is commonly used (Oakley et al., 2006; Oblak et al., 2021). Previous studies reported SWR abnormalities (e.g., shorter SWR duration) in 5xFAD (Iaccarino et al., 2016; Caccavano et al., 2020; Prince et al., 2021). However, at least two issues remain to be addressed. First, although exposure to a novel context in rats or wild-type mice can increase SWR frequency (Cheng and Frank, 2008; Eschenko et al., 2008; Karlsson and Frank, 2008; O’Neill et al., 2008; Joo and Frank, 2018), it is unclear if 5xFAD mice exhibit any deficits in such experience-dependent SWRs.

Second, although amyloid pathology impairs inhibitory connections to pyramidal cells in the hippocampus (Caccavano et al., 2020; Prince et al., 2021), the contributions of extrinsic inputs to SWRs remain to be determined. In particular, septal cholinergic signals play a critical role in SWR genesis (Vandecasteele et al., 2014; Zhang et al., 2021). Since the basal forebrain is a vulnerable area in AD (Whitehouse et al., 1981; Schmitz et al., 2016; Hampel et al., 2018) and 5xFAD mice also develop pathology in basal forebrain cholinergic neurons (Yan et al., 2018; Sun et al., 2022), it is important to determine if the hippocampal cholinergic tone can influence SWRs in 5xFAD mice.

In the present study, we address these two issues. First, we report that 5xFAD mice exhibit experience-dependent SWR deficits. Second, by applying simultaneous *in vivo* electrophysiology and fiber photometry (Patel et al., 2020; Byron and Sakata, 2024), we monitor hippocampal local field potentials (LFPs) together with hippocampal cholinergic signals using AchLightG, a recently developed genetically encoded acetylcholine sensor (Kagiampaki et al., 2023). We report that hippocampal cholinergic tone in 5xFAD mice is intact across sleep-wake cycles.

## Materials and Methods

### Animals

All experiments and procedures were performed in accordance with the United Kingdom Animals (Scientific Procedures) Act of 1986 Home Office regulations and approved by the Home Office (PP0688944) and the University of Strathclyde’s Ethical Committee. 5xFAD mice (JAX006554) were bred with wild-type (WT) mice on the C57BL/6J background (>F10). Breeding scheme was either male 5xFAD x female WT or female 5xFAD x male WT depending on batch. All genotyping was performed by Transnetyx using real-time PCR. 5xFAD+ (5xFAD) and 5xFAD-(WT) mice were housed in a 12 h light/dark cycle with ad libitum access to food and water in pairs of same-sex littermates. 12 mice (6 5xFAD, 3 females and 3 males; 6 WT, 3 females and 3 males) aged 3.5 to 7.5 months were used in electrophysiological experiments, while 14 mice (7 5xFAD, 2 females and 5 males; 7 WT, 3 females and 4 males) aged 4.5 to 9 months were used for simultaneous electrophysiology and fiber photometry experiments.

### *In vivo* electrophysiology

#### Surgical procedure

Mice were anesthetized with isoflurane (3-5% for induction, 1-1.5% for maintenance) delivered with 0.8 L/min air flow and placed on a stereotaxic frame (SR-5M-HT, Narishige) with an incisor bar and ear bars. Body temperature was maintained at 37 °C using a feedback temperature controller (50-7221-F, Harvard Bioscience). The skull was exposed, and two skull screws (418-7123, RS Components) were implanted in the front (AP +1.5 mm from bregma, ML ±1 mm) for cortical electroencephalogram (EEG) and two skull screws in the back (AP -2 mm from lambda, ML +2 mm) as ground and backup. Two wires (1,5 cm, 2840/7, ALPHA WIRE) were inserted into the neck muscles for electromyography (EMG). A craniotomy (AP -2 mm, ML +1.5 mm from bregma) was performed for the insertion of a bipolar electrode into the CA1 (-1.5 mm DV). Bipolar electrodes were composed of two stainless steel wires (AISI 302, 0.1mm diameter, FE205850/2, GoodFellow) of approximately 1.5 cm length. All electrodes were secured with KWIK-KAST (World Precision Instruments) and layers of superglue (918-6872, RS-Pro) and dental cement (Kemdent). After surgery, mice were kept in their home cage on a heat map for 1 h to aid recovery. 24 h after surgery, additional analgesia was provided orally in the form of Rimadyl (20 mg/kg, 0.01% diluted in saline). Mice were single-housed in a high-roofed cage with ad libitum access to water and food. Their recovery was assessed daily for 4 days.

#### Novel environment exposure

After recovery, mice were habituated to handling for 5 consecutive days, followed by 2 weeks of electrophysiological and behavioral testing. In week 1, while mice were placed in a white open field box (30 cm width x 30 cm depth x 40 cm height), EEG and EMG signals were monitored (see below). Each session lasted 2.5 hours. The same procedure was repeated for up to 4 consecutive days. Since the sample size on the fourth day was lower due to recording noise in some animals, the data on the fourth day was excluded.

In week 2, EEG and EMG signals were recorded in their home cages over 5 days. The first day (Day 0) consisted of a 1.5 h home cage recording session followed by 10 min exploration of the open field box used in week 1, which now contained visual landmarks (A4 papers with black stripes and black dots on opposing walls). Days 1 to 4 consisted of a 1 h baseline home cage recording, a 10 min exploration of two identical objects (50 mL falcon tubes filled with sand) in the open field box, followed by a 1.5 h home cage recording. On the last day (Day 4), two additional sessions were included. After the second home cage session, mice were exposed to the open field box with one familiar and one novel object (LEGO blocks) for 10 minutes. This was followed by a third 1.5 h home cage recording. To allow mice to explore the behavioral environment freely, no electrophysiological recording was performed across all behavioral sessions. The open field box and objects were cleaned with 70% ethanol between each session. The entire procedure was repeated four weeks later using a new set of objects (LEGO blocks decorated with tape and polystyrene cylinder decorated with black stars). For data analysis, all data was pooled together.

#### Electrophysiological recording procedure

Electrophysiological EEG and EMG signals were amplified x1000 with a head-stage (HST/32V-G20, Plexon) and main amplifier (PBX3, Plexon). The amplified signals were digitized at 1 kHz using a data acquisition device (NI USB-6211 DAQ, National Instruments) and recorded via a custom-written LabVIEW program (National Instruments). Top-view behavioral videos were also recorded using a webcam (nulaxy, HD 1080p Webcam) at 20 Hz using an additional custom LabVIEW program.

### Simultaneous *in vivo* electrophysiology and fiber photometry

#### Surgical procedure

Mice were anesthetized as described above. A small burr hole was created at AP -2 mm, ML +1.5 mm from bregma for viral injection. A glass micropipette, mounted on a motorized injector (Nanoliter2010, WPI), was slowly inserted into the brain at the first injection site at a depth of 1.65 mm DV, then retracted to the second injection site at 1.4 mm DV. At each depth, 250 nL of viral vectors (AAV2/9-hSyn-AchLightG, titer 1.0 x 10^13^ vg/mL, obtained from the Viral Vector Facility of the University of Zürich) (Kagiampaki et al., 2023) was injected at a rate of 25 nL/min. Two weeks after the viral injection, an implantation surgery was conducted as described above. After the insertion of EEG screws and EMG wires, an optrode was inserted into the hippocampus at AP-2 mm from the bregma, ML -1.5 mm with the tip of the optic fiber at -1.4 mm DV and the tip of the bipolar electrode at -1.5 mm DV. The optrode consisted of a bipolar electrode and an optic fiber (FP400URT, Thorlabs; CF440-10, Thorlabs). The tip of the bipolar electrode was offset from the tip of the fiber by 1 mm and subsequently held in place with superglue (918-6872, RS PRO). After the insertion, skull screws and the optrode were secured with dental cement (Kemdent). Post-surgery care was performed as described earlier.

#### Recording procedures

On Day 0, mice underwent a 2 h mock-recording session to habituate the animals to a tethered condition in their home cage. The recording setup remained off to avoid photobleaching. The next day (Day 1), mice underwent a 1 h baseline recording in their home cage followed by 10 minutes of exploration of two identical objects in the open field box. They also underwent a 1.5 h post-exposure recording session in the home cage.

The fiber photometry setup was described elsewhere (Patel et al., 2020). A 470 nm LED (LED M470L3, Thorlabs) was used for AchLightG excitation, while a 405 nm LED (LED M405L3, Thorlabs) was used to capture acetylcholine-independent isosbestic signals. Light stimulation was controlled via custom-written LabVIEW code at a pulse frequency of 400 Hz with a 50% duty cycle. Synchronization pulses were also generated and saved for offline analysis. The voltage for each LED was adjusted for each recording according to light output measurements for each fiber (1.13-1.48 V for 405 nm LED, 0.87-1.24 V for 470 nm LED) to achieve a target power of 35 μW (0.27 mW/mm^2^). Emission signals were detected via a photodetector (NewFocus 2151, Newport) and digitized together with electrophysiology signals recorded with the same system as described above. All signals were sampled at 5kHz using a data acquisition device (NI USB-6211 DAQ, National Instruments).

### Histology

After the final recording session, mice were deeply anesthetized with pentobarbital and lidocaine and perfused transcardially with phosphate buffered saline (PBS) followed by 4% paraformaldehyde/0.1 M phosphate buffer. Brains were removed and kept in the same fixative overnight at 4°C before then being stored in 30% sucrose PBS for at least 2 days. Brains were cut into 50 μm coronal sections using a sliding microtome (SM2010R, Leica) and stained with DAPI (1:500, Sigma-Aldrich) and Thioflavin-S (0.01%, T1892, Sigma-Aldrich).

Brains of mice that underwent viral injections were cut at 80 µm thickness, washed with PBS and incubated in a blocking solution consisting of 10% Normal Goat Serum (NGS, Thermo Fischer Scientific) in PBST (PBS + 0.3% Triton X-100) for 1 hour at room temperature on a shaker. Sections were then incubated over night at 4°C with mouse monoclonal anti-GFP primary antibody (1:2000, ab1218, Abcam) diluted in 3% NGS in PBST. The next day, sections were washed three times for 5 minutes in PBS, then incubated for 2 hours at room temperature with secondary antibodies (1:1000, goat anti-mouse IgG (H+L) Alexa Fluor, Thermo Fisher Scientific). Sections were washed three times for 5 minutes with PBS and then incubated with DAPI (1:1000, Sigma-Aldrich) and Thiazine Red (0.05%, VWR Chemicals) for 15 minutes. The sections were mounted on gelatin-coated slides and cover-slipped with fluoromount-G (Thermo Fisher Scientific). Then they were imaged under an epifluorescent microscope (Eclipse E600, Nikon) using a custom-written image acquisition program (LabVIEW, National Instruments).

### Data analysis

#### Sleep scoring

Arousal states were classified offline as awake, NREM and REM sleep in 4-s intervals based on EEG and EMG signals using a custom-written MATLAB GUI (Patel et al., 2020; Tsunematsu et al., 2020). Episodes with high EMG power were scored as awake, episodes with low EMG signal and high delta or sigma power were scored as NREM sleep, and episodes with low EMG signal and high theta power were scored as REM sleep.

#### SWR detection

The procedure was essentially the same as described elsewhere (Tsunematsu et al., 2020; Tsunematsu et al., 2023). Hippocampal LFP signals were bandpass filtered at 140-250 Hz with a 3rd-order Butterworth filter and the envelope of the filtered signal was obtained by computing the root-mean-square (RMS) over a sliding window of 20 ms. Signals during REM sleep episodes were used to compute two thresholds: the higher threshold was calculated as mean signal + 5 x standard deviation (SD) and the lower threshold was calculated as mean signal + 2 x SD. For recordings without REM sleep episodes, a 12-30-s noise-free NREM sleep segment, typically occurring at the end of a NREM sleep episode, was manually selected and used for threshold estimation. SWR candidates were identified as events where the signal power exceeded the higher threshold. Event onset and offset were defined as time points at which the signal power crossed the lower threshold. Overlapping events were merged and events with a duration shorter than 20 ms or longer than 300 ms were removed. Artefacts caused by muscle twitching during sleep were excluded by comparing the EMG RMS values during each SWR candidate to a threshold defined as twice EMG RMS values during REM episodes or noise-free NREM sleep segments. SWR candidate events exceeding the threshold were classified as artefacts and excluded from analysis.

#### Sleep spindle detection

Sleep spindle detection was the same as in a previous study (Copping and Silverman, 2021). EEG signals were bandpass filtered using a custom-designed Butterworth filter applying a first stopband frequency at 3 Hz, first passband frequency at 10 Hz, second passband frequency at 15 Hz, second stopband frequency at 22 Hz and a stopband attenuation level of 24 dB. The envelope of the filtered signal was obtained by computing the RMS over a sliding window of 750 ms. A high threshold (mean x 3.5) and a low threshold (mean x 2) were calculated based on NREM sleep signals. Spindle candidates were identified when the signal power exceeded the higher threshold. The onset and offset were determined based on the lower threshold. Events with <0.5 s or >10 s were excluded. Lastly, events with high EMG power were excluded based on the same EMG threshold method as described above.

#### SWR–spindle coupling

To assess the coupling between SWRs and sleep spindles, hippocampal LFP signals were bandpass filtered between 140-250 Hz using a third-order Butterworth filter. The signal envelope was obtained by computing the RMS over a 20-ms sliding window. Spindle onset times, detected in the EEG signal, were used to define 4 s peri-spindle windows (-2 to +2 s from spindle onset). Within these windows, the hippocampal LFP envelope was Z-scored based on the pre-spindle baseline period (-2 to -1 s). The mean Z-scored signal was computed for each recording.

#### Photometry signal processing

Photometry signals were analyzed using a method modified from a previously published protocol (Patel et al., 2020). First, based on the synchronization pulses of the two LEDs, the median signal during each illumination period was computed for both 405 nm and 470 nm excitations. Second, a third-order Butterworth low-pass filter with a 2 Hz cutoff was applied to reduce high frequency noise. Third, photobleaching for each excitation signal was corrected using double-exponential fitting. Fourth, the corrected 405 nm signal was linearly scaled and subtracted from the corrected 470 nm signal to reduce motion artifacts. Finally, the signals were Z-scored by computing the mean and standard deviation of the entire signals.

To correlate AchLightG signals with EEG sigma or hippocampal ripple power, AchLightG signals were bandpass-filtered with 0.01 Hz cutoff whereas EEG sigma and hippocampal ripple power were computed by applying a bandpass filter (10-15 Hz for EEG sigma; 140-250 Hz for hippocampal ripples) and calculating room-mean-square in a 10-s window with 99% overlap. Because of frequent outlier signals, Spearman’s rank correlation was calculated based on signals during NREMS only.

To assess the infraslow components of AchLightG signals, a continuous wavelet transform (*cwt* function in MATLAB) was applied to the entire length of downsampled (10 Hz) signals and extracted the power of three frequency bands: <0.01 Hz, 0.01-0.1 Hz, and 0.1-1 Hz.

#### Behavioral analysis

Videos of mice exploring objects in an open field box were analyzed using DeepLabCut (Mathis et al., 2018). For training, the following body parts were manually marked on 300 randomly selected frames: nose, left ear, right ear, neck, left shoulder, right shoulder, left hip, right hip, and tail base. After a network was trained for 500,000 iterations, videos were processed and videos with markers for all body parts were created to inspect the accuracy of labelling. Coordinates of each body part were then analyzed in MATLAB. The distance between the nose and each object in an open field box was calculated. Exploration behavior was defined as periods in which the nose remained within a defined radius of 2 cm from an object’s outer edge within the first 5 min of recording. The discrimination index was calculated as the difference in exploration time between object A and object B, divided by the total exploration time of both objects.

### Statistical analysis

All data was presented as mean ± SEM unless stated otherwise. The following statistical tests were performed in MATLAB: unpaired t-test, one-way ANOVA, two-way ANOVA and repeated measures ANOVA, depending on data types. A p-value <0.05 was considered significant. Tukey’s honest significant difference (HSD) test was performed for post-hoc comparisons.

## Results

### Experience-dependent increase in SWRs is impaired in 5xFAD mice

To examine if 5xFAD mice exhibit deficits in hippocampal SWRs, animals were placed in an open field while cortical EEGs and hippocampal LFPs were continuously monitored for 2.5 h (**Fig. 1A**). We repeated the same procedure for at least three days. We chronically implanted electrodes around the CA1 pyramidal cell layer (*stratum pyramidale*) (**Fig. 1B and Supplementary Fig. 1A**) to detect SWRs offline (**Fig. 1C**).

**Figure 1.**
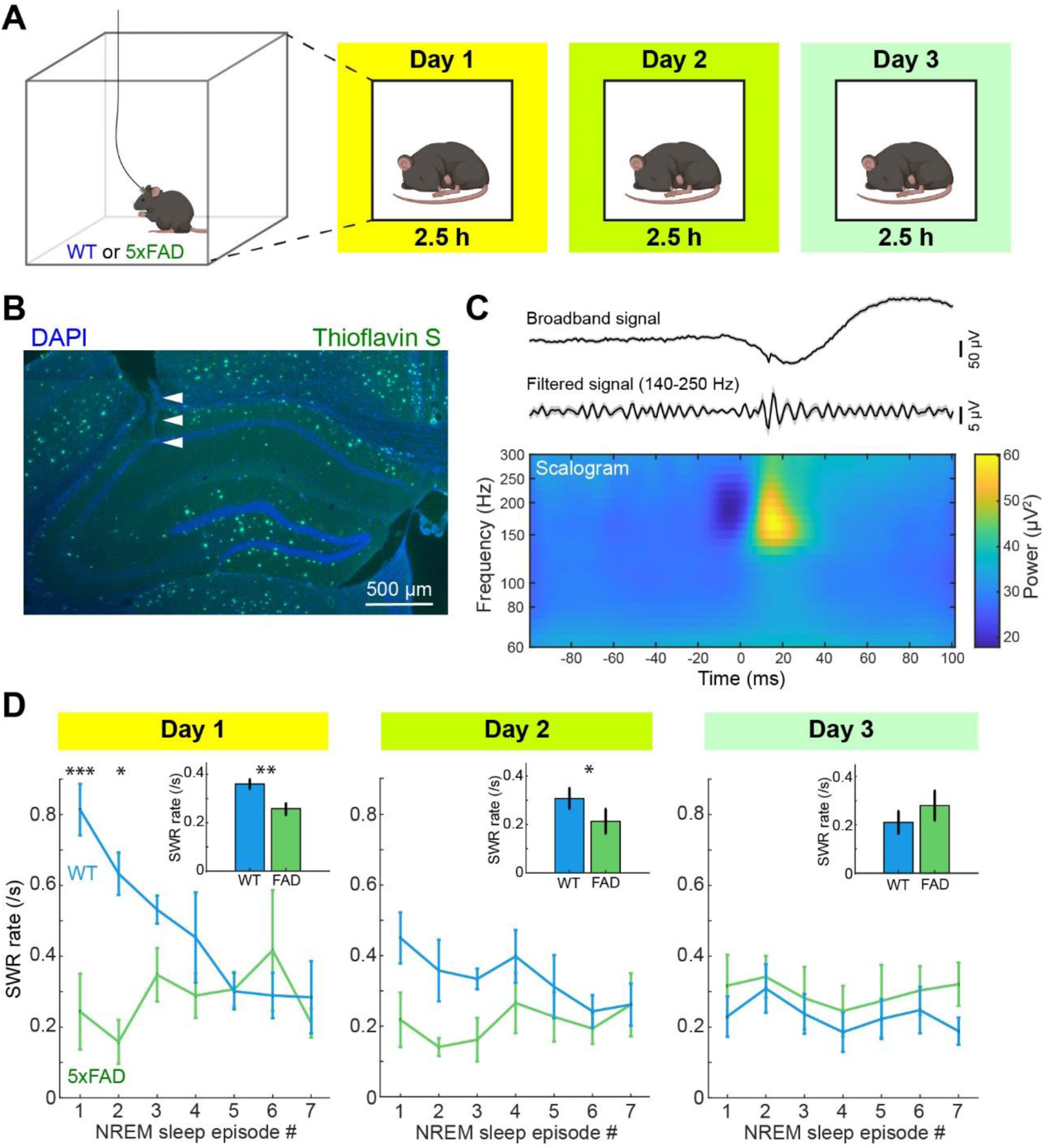
Deficits in transient SWR increase in 5xFAD mice after exposure to a novel environment. **A.** Diagram to outline the experimental design. **B.** A histological image showing the track of an electrode on the CA1 pyramidal cell layer (triangles) and amyloid plaques (green). **C.** An example of detected SWRs. Top, average broadband signals showing a sharp wave. Middle, average bandpass filtered signals showing ripples. Bottom, average scalogram of detected SWRs. Time 0 is SWR onset. **D.** Dynamics of SWR rate across NREMS episodes and recording days. The inset indicates the overall SWR rate during NREMS. *, *p* < 0.05; **, *p* < 0.01; ***, *p* < 0.001; two-way ANOVA with post-hoc Tukey’s HSD test; t-test (inset).

Based on this data set (WT, n = 6; 5xFAD, n = 6), we asked if the frequency of SWRs changed across NREMS episodes while the animals were in the open field (**Fig. 1D**). On Day 1, we observed that the SWR rate was high at early NREMS episodes and decreased gradually in WT mice. On the other hand, 5xFAD mice did not exhibit such a transient increase in the SWR rate, leading to a significantly lower rate in the first two NREMS episodes in 5xFAD mice compared to WT mice (F(6, 70) = 4.34, *p* < 0.001, two-way ANOVA with post-hoc Tukey’s HSD test). As a result, the overall SWR rate across NREMS episodes was significantly lower in 5xFAD mice (*p* = 0.002, t-test). A similar trend was observed on Day 2, but not on Day 3. The average duration of NREMS episodes was comparable between animal groups (**Supplementary Fig. 1B**). Thus, 5xFAD mice exhibited deficits in a transient SWR increase during NREMS after exposure to a novel environment.

### No experience-dependent changes in sleep spindles in both WT and 5xFAD mice

Since sleep spindle is another electrophysiological signature of NREMS and has been implicated in AD (Rasch and Born, 2013; Weng et al., 2020), we also examined if sleep spindles also increased after exposure to a novel environment in WT mice, and if 5xFAD mice showed any abnormalities in sleep spindles (**Fig. 2**). We detected sleep spindles during NREMS by processing cortical EEG signals (**Fig. 2A**). The rate of sleep spindles was generally low (0.01 - 0.05 Hz). We did not observe any significant differences in spindle rate between animal groups across days (*p* = 0.11 on Day 1; *p* = 0.67 on Day 2; *p* = 0.65 on Day 3; t-test) (**Fig. 2B**). Thus, although SWRs exhibited an experience-dependent transient increase in WT mice, sleep spindles were insensitive to experience in our experimental condition.

**Figure 2.**
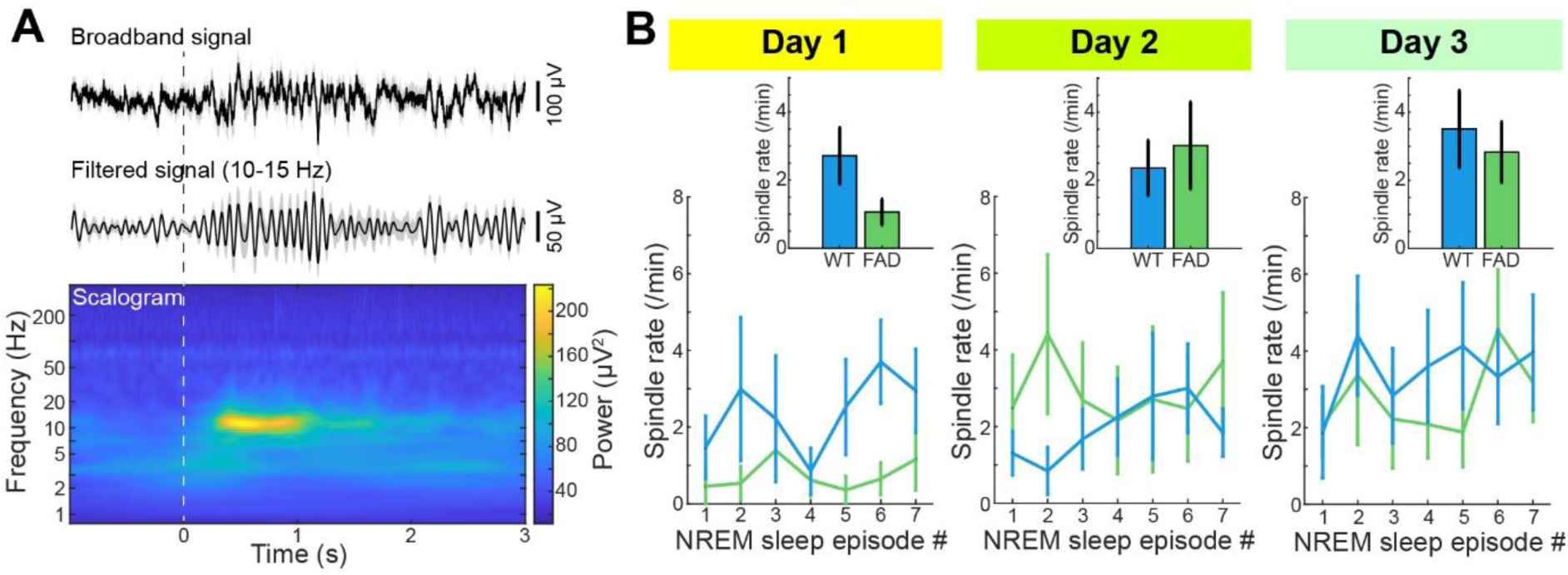
Sleep spindles and their experience dependence in WT and 5xFAD mice. **A.** An example of detected sleep spindles. Top, average broadband signals. Middle, average bandpass filtered signals showing sleep spindles. Bottom, average scalogram of detected sleep spindles. Time 0 is spindle onset. **B.** Dynamics of spindle rate across NREMS episodes and recording days. The inset indicates the overall spindle rate during NREMS. Error bars indicate SEM.

### Novel objects increase SWR rate in WT, but not in 5xFAD mice

In the following weeks, we further examined if after exposure to a different environment, we could observe a transient increase in SWRs in WT mice and a similar deficit in 5xFAD mice (**Fig. 3**). We used the same open field but placed two objects at the corners of the arena, which we called a behavioral session (**Fig. 3A**). We repeatedly exposed animals to this condition for four days. On Day 4, we introduced a second behavioral session by replacing one of the objects. Electrophysiological recording was performed in their home cage for 1.5 h whereas animals were allowed to explore the arena and objects without being tethered.

**Figure 3.**
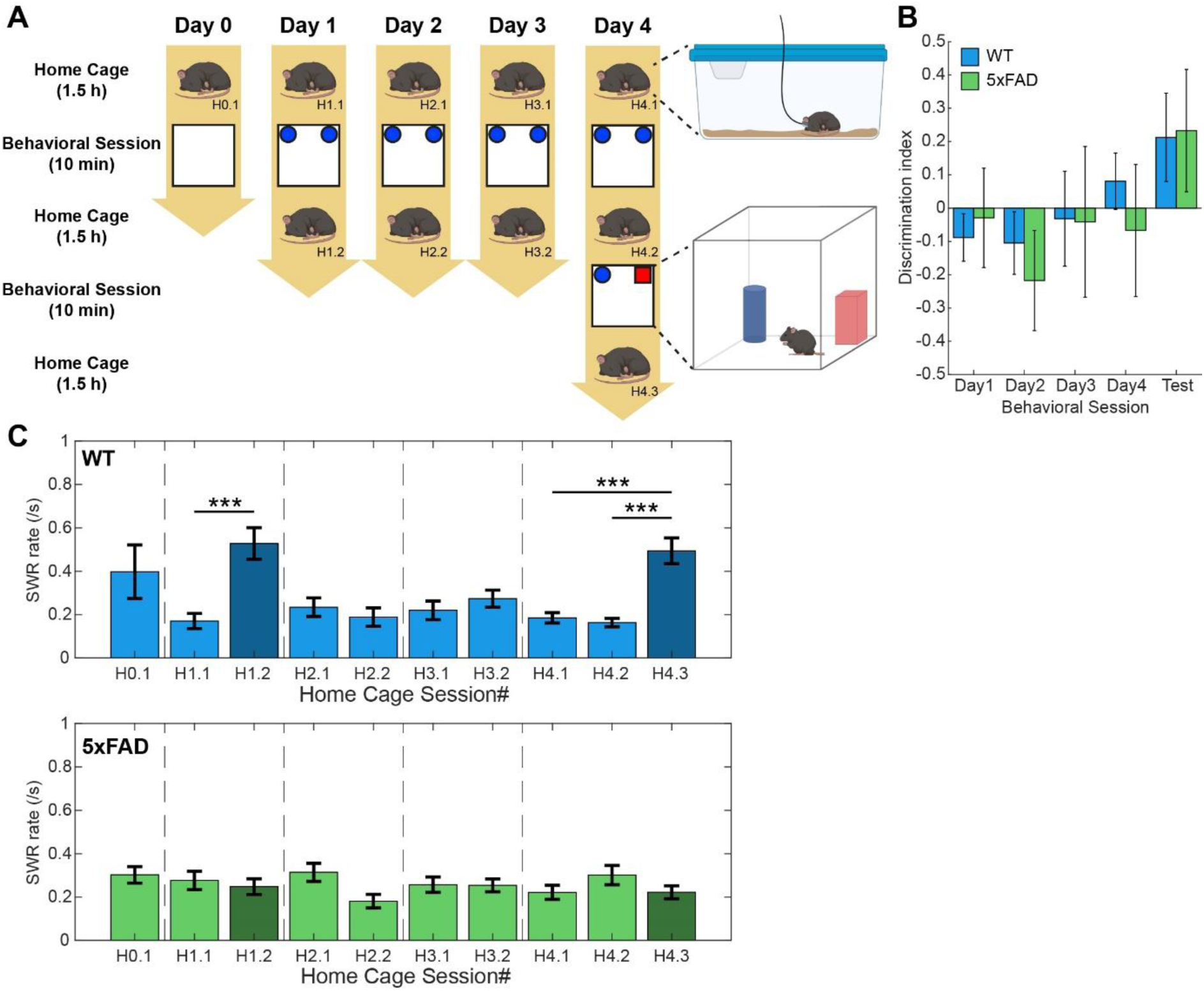
Novel objects induce SWR increase in WT, but not in 5xFAD mice. **A.** Experimental design. Electrophysiological recording was performed during the home cage session whereas animals were allowed to explore their environment freely during the behavioral session. **B.** Discrimination index across behavioral sessions. **C.** SWR rate during NREMS across home cage sessions. The annotation number indicates Day and home cage session on the same day. For example, H0.1 means the first home cage session on Day 0. ***, *p* < 0.0001; one-way ANOVA with post-hoc Tukey HSD test. The significance was assessed within the same recording day (e.g., H1.1 vs H1.2).

First, we assessed the animal’s exploratory behavior toward the objects by tracking their nose. To quantify their discriminatory behavior between two objects, we computed the discrimination index: as an animal explores the right-side object longer, the index becomes positive and increases. When a novel object was introduced, both WT and 5xFAD mice tended to explore it more (**Fig. 3B**). However, the effect was small and not statistically significant (F(1,19) = 0.55, *p* = 0.47, repeated measures ANOVA).

Although we did not find any noticeable behavioral changes across the behavioral sessions, we observed a transient increase in SWRs during NREMS in WT mice when the environment was modified (**Fig. 3C**). More specifically, two objects were introduced on Day 1 and a novel object was introduced in the second behavioral session on Day 4. In the subsequent home cage sessions, we observed a significant increase in SWR rate (F(9,90) = 5.65, *p* < 0.001, one-way ANOVA with post-hoc Tukey’s HSD test). However, we did not observe such an increase in 5xFAD mice (F(9,100) = 1.33, *p* = 0.23, one-way ANOVA). These results strengthened the initial observation in **Figure 1**.

### Intact spindle-SWR coupling in 5xFAD mice

Similar to the initial experiment (**Fig. 2**), we also assessed sleep spindles across home cage sessions (**Fig. 4A**). The overall spindle rate was comparable to the initial experiment, and we did not observe any significant changes across sessions in both animal groups (WT: F(9,90) = 0.74, *p* = 0.67, 5xFAD: F(9,100) = 0.46, *p* = 0.90, one-way ANOVA).

**Figure 4.**
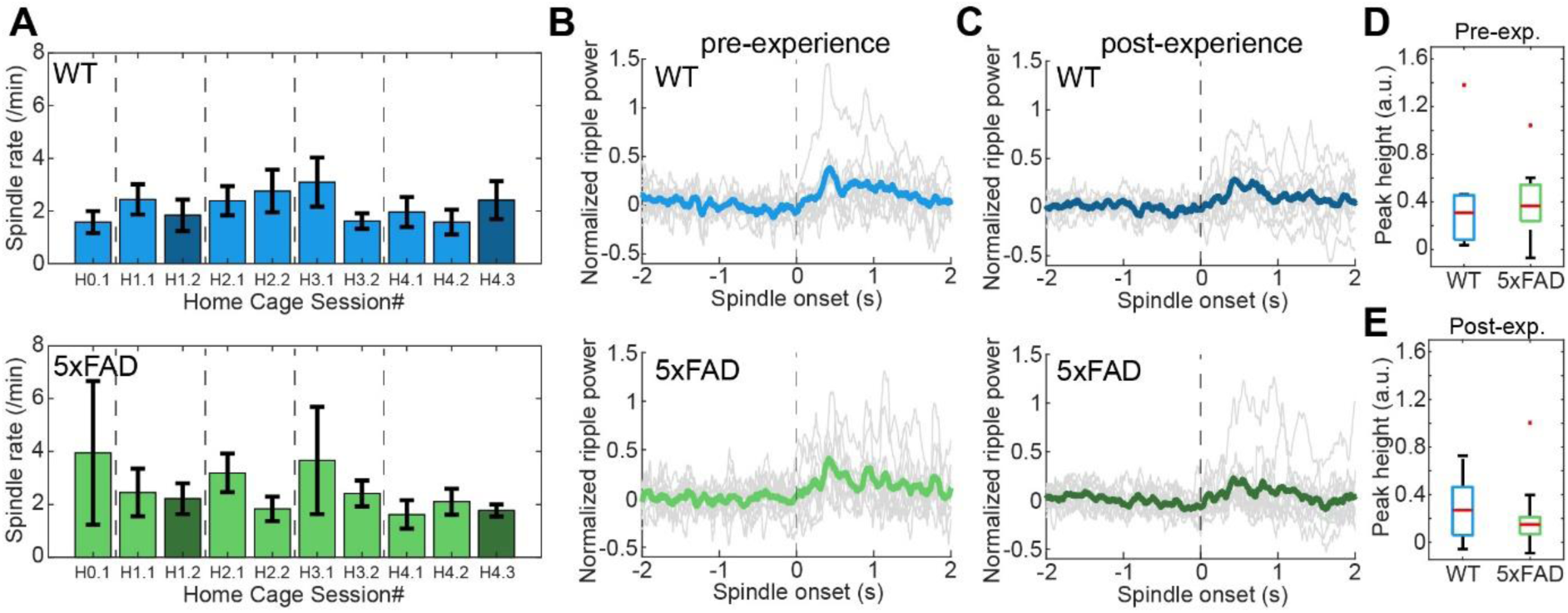
Effects of novel objects on sleep spindle rate and SWR-spindle coupling. **A.** Sleep spindle rate across home cage sessions in WT (*top*) and 5xFAD mice (*bottom*). The dark blue and green indicate sessions after novel objects were introduced. **B.** Normalized (z-scored) sleep spindle power triggered by the onset of SWRs in WT (top) and 5xFAD mice (bottom) during home cage sessions before introducing novel objects (pre-experience sessions). Gray lines indicate the average profile of each session. The thick line indicates the overall average profile. **C.** Normalized (z-scored) sleep spindle power triggered by the onset of SWRs in WT (top) and 5xFAD mice (bottom) during home cage sessions after introducing novel objects (post-experience sessions). **D.** The peak height of normalized spindle power triggered by the onset of SWRs during home cage sessions before introducing novel objects (pre-experience session). **E.** The peak height of normalized spindle power triggered by the onset of SWRs during home cage sessions after introducing novel objects (post-experience session).

Since the functional coupling between sleep spindles and SWRs has been implicated in sleep-dependent memory consolidation (Staresina et al., 2015; Klinzing et al., 2019; Staresina et al., 2023), we examined if a similar functional coupling could be seen in our experimental condition and if the coupling differed between animal groups (**Figs. 4B-E**). To investigate the coupling between SWR onset and sleep spindle power, we separated home cage sessions into “pre-experience” or “post-experience” sessions depending on whether novel objects were introduced (sessions H1.2 and H4.3) or not. We consistently observed a transient increase in spindle power after SWRs in both types of sessions and for both animal groups (**Figs. 4B and C**). To compare the coupling between animal groups, we calculated the peak height of normalized spindle power after SWR onset (**Figs. 4D and E**). This value was comparable between animal groups in the pre-experience session (*p* = 0.93, t-test) (**Fig. 4D**) and the post-experience session (*p* = 0.52, t-test) (**Fig. 4E**). Thus, although SWRs transiently increased after introducing novel objects only in WT mice, sleep spindles and the coupling between SWRs and sleep spindles were stable across sessions and animal groups.

### Simultaneous monitoring of hippocampal cholinergic and electrophysiological signals across sleep-wake cycles

To explain the deficit in experience-dependent SWR increase in 5xFAD mice, we hypothesized that hippocampal cholinergic signals play a role because septal cholinergic inputs inhibit SWRs (Vandecasteele et al., 2014; Zhang et al., 2021) and amyloid pathology can be seen in the basal forebrain in 5xFAD mice (Yan et al., 2018; Sun et al., 2022). To test this hypothesis, we conducted simultaneous LFP recording and fiber photometry in WT (n = 7) and 5xFAD mice (n = 7) (**Fig. 5A**). To monitor cholinergic signals, we expressed a genetically encoded acetylcholine sensor, AchLightG (Kagiampaki et al., 2023), in the hippocampus (**Fig. 5C**). To minimize photobleaching effects, we modified the experimental design, in which photometry experiments were conducted during home cage sessions before and after a behavioral session only on Day 1 (**Fig. 5B**). During a home cage session, we observed state-dependent changes in cholinergic signals along with spectral changes in hippocampal LFP signals across sleep-wake cycles (**Fig. 5D and Supplementary Fig. 2**).

**Figure 5.**
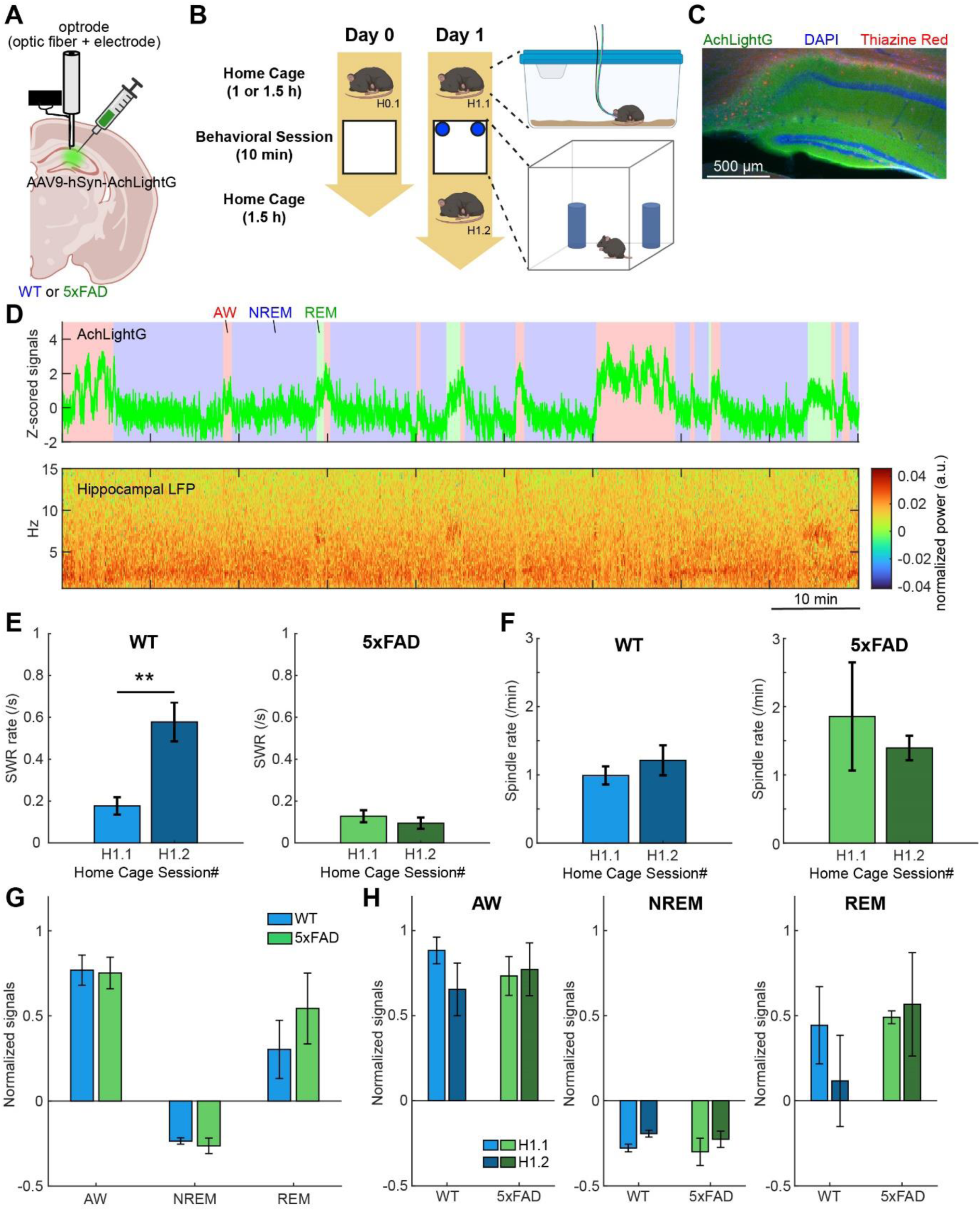
State-dependent cholinergic signals in the hippocampus of WT and 5xFAD mice. **A.** Schematic drawing of the experimental configuration. **B.** Experimental design. **C.** An image showing AchLightG (green) and Thiazine-Red-stained amyloid plaques (red). DAPI signals outline the cytoarchitecture in the hippocampus. **D.** An example of an AchLightG trace across sleep-wake states (*top*) and a spectrogram of hippocampal LFP signals (*bottom*) in a WT mouse. **E.** SWR rate during NREMS in home cage sessions in WT (*left*) and 5xFAD mice (*right*). **, *p* < 0.005, t-test. **F.** Sleep spindle rate during NREMS in home cage sessions in WT (*left*) and 5xFAD mice (*right*). **G.** The average normalized (Z-scored) AchLightG signal across three states in WT and 5xFAD mice. AW, wakefulness; NREM, NREM sleep; REM, REM sleep. **H.** The average normalized (Z-scored) AchLightG signals across individual home cage sessions.

First, we re-confirmed that the SWR rate increased during NREMS after introducing objects in WT mice (*p* < 0.005, t-test), but not in 5xFAD mice (*p* = 0.14, t-test) (**Fig. 5E**). On the other hand, the sleep spindle rate did not change (**Fig. 5F**). Second, we compared the average fluorescent signals between animal groups across arousal states (**Fig. 5G**) and home cage sessions (**Fig. 5H**). While cholinergic signals were lower during NREMS compared to wakefulness and REM sleep as expected (F(2,72) = 57.1, *p* < 0.001, two-way ANOVA for the effect of vigilance states), we did not observe any differences between animal groups (F(2,72) = 0.86, *p* =0.42, two-way ANOVA for interaction between animal groups and vigilance states) (**Fig. 5G**). This was also the case when we split the data into two home cage sessions (H1.1 and H1.2) (AW, F(1,27) = 1.08, *p* = 0.30; NREM, F(1,27) = 0.01, *p* = 0.91; REM, F(1,16) = 0.4, *p* = 0.53, one-way ANOVA) (**Fig. 5H**). Thus, the gross cholinergic tone in the hippocampus was comparable in both animal groups.

### Intact hippocampal cholinergic dynamics during NREM sleep in 5xFAD mice

Next, we examined if hippocampal cholinergic dynamics during NREMS could explain the SWR deficit in 5xFAD mice. To this end, we first correlated cholinergic dynamics with hippocampal ripple and cortical sigma power during NREMS (**Figs. 6A-C**). Although hippocampal cholinergic tones were generally low during NREMS as described above (**Fig. 5**), we noticed that hippocampal cholinergic tones fluctuated slowly. To characterize this trend, we applied a lowpass (<0.01 Hz) filter and computed the corresponding hippocampal ripple (140-250 Hz) and cortical sigma (10-15 Hz) power (**Fig. 6A**). Interestingly, the infraslow dynamics of cholinergic tones were negatively correlated with both electrophysiological signals, especially with cortical sigma power during NREMS (**Fig. 6A**). We examined if this correlation was changed by experience and if 5xFAD mice exhibited any differences, with respect to ripple (**Fig. 6B**) and sigma power (**Fig. 6C**). Although we confirmed the non-trivial negative correlation between cholinergic tones and cortical sigma power consistently, we did not observe any differences between conditions and animal groups (vs HC ripple, F(1,27) = 0.04, *p* = 0.84; vs EEG sigma, F(1,27) = 1.07, *p* = 0.31, two-way ANOVA) (**Figs. 6B and C**).

**Figure 6.**
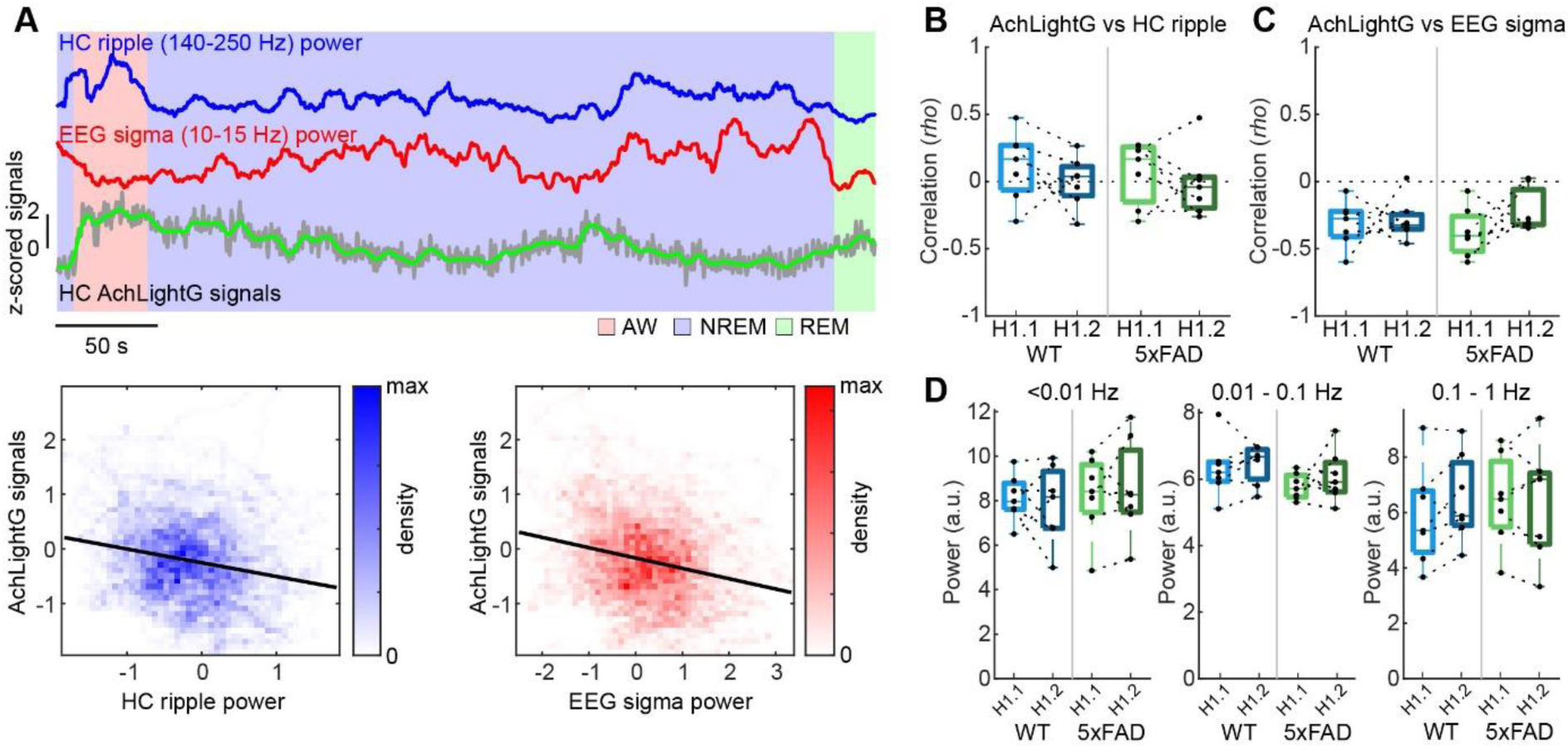
Hippocampal cholinergic dynamics during NREM sleep in WT and 5xFAD mice. **A.** *top*, an example of infraslow fluctuations in hippocampal (HC) ripple (blue), cortical sigma (red) and hippocampal cholinergic signals (green). *Bottom*, an example of scatter plots showing the negative correlation of AchLightG signals with HC ripple power (*left*, blue) and cortical sigma power (*right*, red). Data points were extracted only from NREM sleep episodes and presented as a density map. The solid line indicates a linear regression line. **B.** Spearman’s correlation *rho* between AchLightG and HC ripple signals during NREMS across home cage sessions and animal groups. **C.** Spearman’s correlation *rho* between AchLightG and cortical sigma signals during NREMS across home cage sessions and animal groups. **D.** Spectral components of AchLightG signals across home cage sessions and animal groups.

Finally, we compared cholinergic dynamics across three frequency bands (<0.01 Hz, 0.01-0.1 Hz, and 0.1-1 Hz) (**Fig. 6D**). None of the band powers showed noticeable differences between home cage conditions or animal groups (<0.01 Hz, F(1,27) = 0.21, *p* = 0.65; 0.01-0.1 Hz, F(1,27) = 0.11, *p* = 0.74; 0.1-1 Hz, F(1,27) = 0.29, *p* = 0.59). Overall, these results suggest that hippocampal cholinergic signals cannot explain SWR deficits in 5xFAD mice.

## Discussion

In the present study, we investigated whether 5xFAD mice exhibit any deficits in experience-dependent SWRs. If so, we investigated how hippocampal cholinergic tones play a role. While we consistently observed the impairment of an experience-dependent increase in SWRs during NREMS across all three experimental conditions (**Figs. 1, 3 and 5**), sleep spindles were intact (**Figs. 2, 4 and 5**). We also characterized hippocampal cholinergic tones across sleep-wake cycles and the relationship between cholinergic and ripple power profiles during NREMS. We found that hippocampal cholinergic signals were generally spared from amyloid pathology (**Figs. 5 and 6**).

### Experience-dependent SWR increase and deficit in 5xFAD mice

An experience-dependent transient increase in SWRs has been reported in rats and mice (Cheng and Frank, 2008; Eschenko et al., 2008; Karlsson and Frank, 2008; O’Neill et al., 2008). Although abnormalities in SWRs were reported in 5xFAD mice (Iaccarino et al., 2016; Caccavano et al., 2020; Prince et al., 2021), we have demonstrated for the first time that 5xFAD mice exhibit deficits in experience-dependent increases in SWR rate. A functional implication of our observations remains unclear as we did not observe significant differences in the performance of the novel object recognition test. However, since experience-dependent SWR increases has been implicated in memory consolidation (Cheng and Frank, 2008; Eschenko et al., 2008), memory retrieval (Joo and Frank, 2018), and metabolic regulation (Tingley et al., 2021; Kaya et al., 2025), 5xFAD mice may show impairments in some aspects.

Both intrinsic and extrinsic components contribute to experience-dependent SWR increases. For example, within the hippocampus, parvalbumin-positive (PV+) interneurons regulate ripple frequency and the synchronous firing of pyramidal cells (Schlingloff et al., 2014; Buzsaki, 2015; Ognjanovski et al., 2017; Vancura et al., 2023). Cholecytokinin-positive (CCK+) interneurons also modulate SWRs (Vancura et al., 2023). In 5xFAD mice, abnormalities in hippocampal PV+ cells are well documented. As an extrinsic component, although septal cholinergic signals onto the hippocampus also play a causal role in SWR generation (Vandecasteele et al., 2014; Zhang et al., 2021), no previous studies investigated hippocampal cholinergic tones in 5xFAD mice. Therefore, we monitored hippocampal cholinergic signals across sleep-wake cycles by utilizing a novel sensor, AchLightG, while simultaneously monitoring hippocampal LFPs in 5xFAD mice for the first time. However, we did not identify any significant differences in hippocampal cholinergic signals between the animal groups. These results support the notion that intrinsic components may better explain the SWR deficit in 5xFAD mice. Further studies are needed to determine the mechanism.

### Sleep spindles in AD mice

Several studies reported a decrease in sleep spindles in several AD mouse models (Benthem et al., 2020; Altunkaya et al., 2024; Campbell et al., 2024) whereas another reported an increase in sigma power (Filon et al., 2020) and one study found no changes after 40-Hz sensory stimulation (Cushing et al., 2024). In the present study, we did not identify any significant differences in sleep spindle rate between WT and 5xFAD mice across all conditions. The discrepancy between the studies may stem from various factors, such as experimental and analytical parameters. Since plaque distribution varies across thalamic regions in 5xFAD mice (Oakley et al., 2006; Oblak et al., 2021), it is possible that sleep spindles are differently affected across cortical regions.

### Infraslow dynamics of hippocampal cholinergic signals and sigma power

Unexpectedly, we discovered a negative correlation between hippocampal cholinergic dynamics and cortical sigma power during NREMS regardless of genotype. Infraslow fluctuations of noradrenaline during NREMS have been recently reported (Osorio-Forero et al., 2025) and microarousals and the activity of the glymphatic system are associated with these infraslow dynamics (Hauglund et al., 2025; Luthi and Nedergaard, 2025). Our findings complement this emerging topic by demonstrating that hippocampal cholinergic signals also co-fluctuate with cortical sigma power in an infraslow range during NREMS. It would be interesting in the future to investigate further how various components (e.g., cortical sigma power, cholinergic signals, and other neuromodulatory signals – dopamine and norepinephrine) are coordinated globally, how they influence hippocampal functions, and how such global coordination and the activity of the glymphatic system are disrupted in AD.

### Limitations of the study

There are at least three limitations in this study that need to be addressed in the future. First, although we consistently observed impaired SWRs in 5xFAD mice, the relevance to behavior or cognitive functions remains to be explored. In addition to utilizing various different behavioral tasks, combining them with an advanced analytical approach (Sakata, 2023; Tillmann et al., 2024; Weinreb et al., 2024) may be a promising avenue to address this issue.

Second, since we monitored only LFPs, monitoring individual neural activity in the hippocampus will provide invaluable information in the future. Although we did not observe significant behavioral changes between animal groups, it is possible that place fields of hippocampal populations may exhibit detectable differences between animal groups.

Finally, while we characterized hippocampal cholinergic signals, the exact mechanism of SWR deficits in 5xFAD mice remains to be determined. Since disrupted somatic inhibition in the hippocampus has been well documented (Caccavano et al., 2020; Prince et al., 2021), it would be interesting to investigate experience-dependent changes in PV+ neuron activity between WT and 5xFAD mice.

### Conclusions

5xFAD mice exhibited impairment of experience-dependent SWR increase. Even though hippocampal cholinergic tones play a critical role in SWR generation, cholinergic signals are generally spared in 5xFAD mice. This implies that amyloid pathology affects non-cholinergic components to disrupt experience-dependent SWR increases.

## Authors Contributions

- Conceptualization: PS and SS
- Methodology: PS, NB, SG and SS
- Software: PS and SS
- Validation: PS and SS
- Formal analysis: PS and SS
- Investigation: PS, SG, and APC
- Resources: VR, TP and SS
- Data Curation: PS and SS
- Writing – Original Draft: PS and SS
- Writing – Review & Editing: PS, NB, SG, VR, TP and SS
- Visualization: PS and SS
- Supervision: SS
- Project administration: SS
- Funding acquisition: TP and SS

## Acknowledgements

This work was supported by Medical Research Council (MR/V033964/1 and MR/Y004051/1 to S.S.), and the European Union’s Horizon 2020 (ERC Starting Grant 891959 to T.P., and H2020-ICT, DEEPER, 101016787 to T.P. and S.S).

