## Supplementary Figures for "Experience-dependent sharp-wave ripple deficits in an Alzheimer’s disease mouse model"

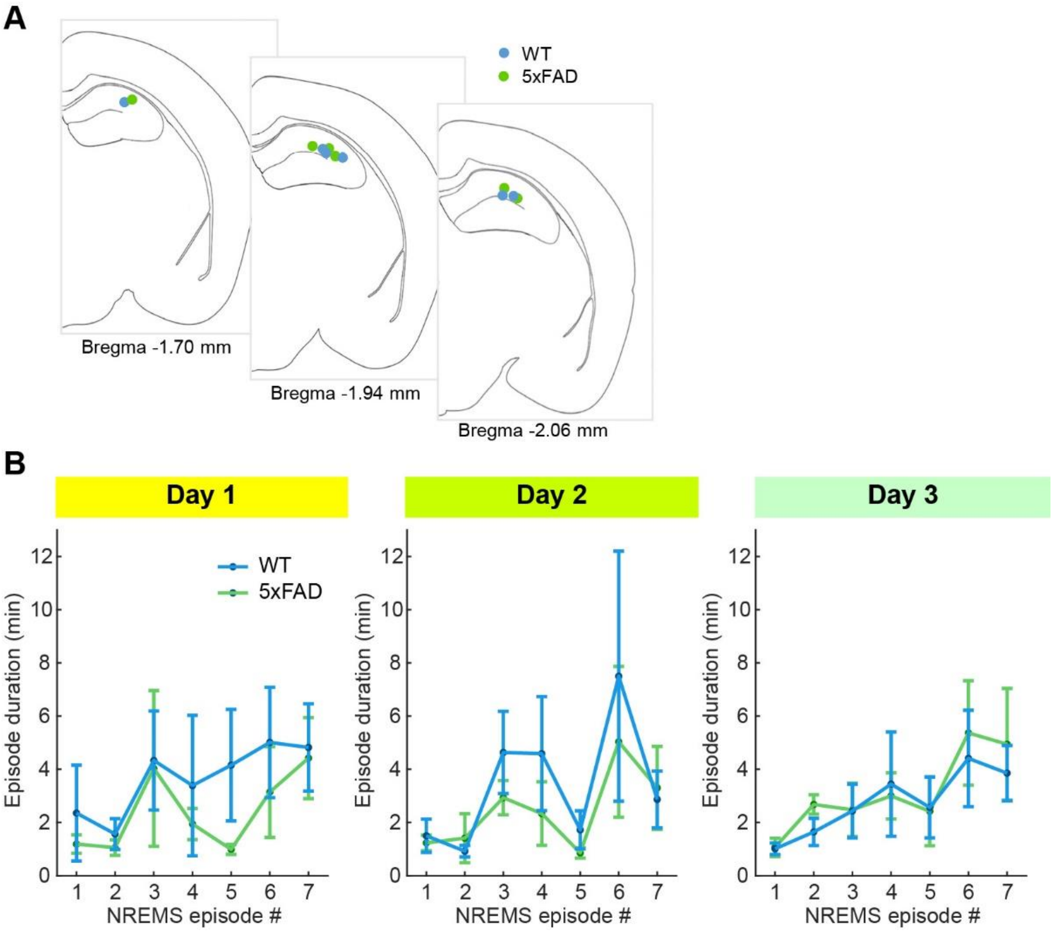


**Supplementary Figure 1. Electrode tip positions and NREMS episode duration.**

**A**. Drawing of electrode tip positions in WT and 5xFAD mice.

**B**. The average duration of NREMS episodes. Day 1: $F(6,70) = 0.18$, $p = 0.98$. Day 2: $F(6,70) = 0.24$, $p = 0.96$. Day 3: $F(6,70) = 0.12$, $p = 0.99$. Two-way ANOVA.

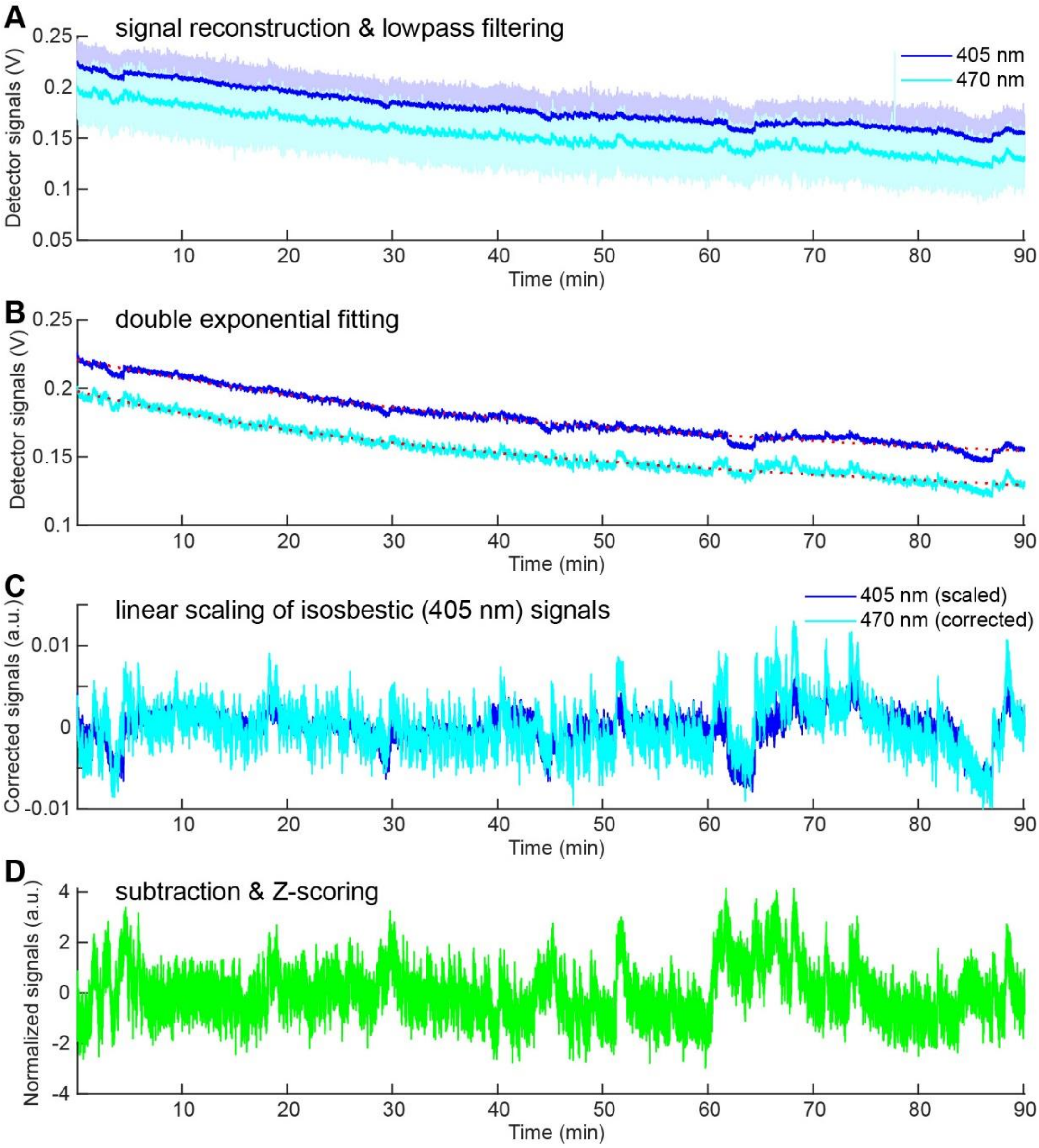


**Supplementary Figure 2. AchLightG signal processing workflow.**

**A**. Since 470 and 405 nm pulses were alternated, photodetector signals for each wavelength were analyzed separately by computing the median of signals for each pulse (signal reconstruction). The reconstructed signals were lowpass-filtered (cut-off frequency, 2 Hz).

**B**. The filtered signals were fitted with a double exponential curve to estimate a photobleaching profile (red dotted line).

**C**. Corrected signals for 405 nm pulses were linearly scaled to estimate acetylcholine-independent artifacts (e.g., motion artifacts).

**D**. The subtracted signals were Z-scored.
